# Shared Genetic Risk between Eating Disorder- and Substance-Use-Related Phenotypes: Evidence from Genome-Wide Association Studies

**DOI:** 10.1101/741512

**Authors:** Melissa A. Munn-Chernoff, Emma C. Johnson, Yi-Ling Chou, Jonathan R.I. Coleman, Laura M. Thornton, Raymond K. Walters, Zeynep Yilmaz, Jessica H. Baker, Christopher Hübel, Scott Gordon, Sarah E. Medland, Hunna J. Watson, Héléna A. Gaspar, Julien Bryois, Anke Hinney, Virpi M. Leppä, Manuel Mattheisen, Stephan Ripke, Shuyang Yao, Paola Giusti-Rodríguez, Ken B. Hanscombe, Roger A.H. Adan, Lars Alfredsson, Tetsuya Ando, Ole A. Andreassen, Wade H. Berrettini, Ilka Boehm, Claudette Boni, Vesna Boraska Perica, Katharina Buehren, Roland Burghardt, Matteo Cassina, Sven Cichon, Maurizio Clementi, Roger D. Cone, Philippe Courtet, Scott Crow, James J. Crowley, Unna N. Danner, Oliver S.P. Davis, Martina de Zwaan, George Dedoussis, Daniela Degortes, Janiece E. DeSocio, Danielle M. Dick, Dimitris Dikeos, Christian Dina, Monika Dmitrzak-Weglarz, Elisa Docampo, Laramie E. Duncan, Karin Egberts, Stefan Ehrlich, Geòrgia Escaramís, Tõnu Esko, Xavier Estivill, Anne Farmer, Angela Favaro, Fernando Fernández-Aranda, Manfred M. Fichter, Krista Fischer, Manuel Föcker, Lenka Foretova, Andreas J. Forstner, Monica Forzan, Christopher S. Franklin, Steven Gallinger, Ina Giegling, Johanna Giuranna, Fragiskos Gonidakis, Philip Gorwood, Monica Gratacos Mayora, Sébastien Guillaume, Yiran Guo, Hakon Hakonarson, Konstantinos Hatzikotoulas, Joanna Hauser, Johannes Hebebrand, Sietske G. Helder, Stefan Herms, Beate Herpertz-Dahlmann, Wolfgang Herzog, Laura M. Huckins, James I. Hudson, Hartmut Imgart, Hidetoshi Inoko, Vladimir Janout, Susana Jiménez-Murcia, Antonio Julià, Gursharan Kalsi, Deborah Kaminská, Leila Karhunen, Andreas Karwautz, Martien J.H. Kas, James L. Kennedy, Anna Keski-Rahkonen, Kirsty Kiezebrink, Youl-Ri Kim, Kelly L. Klump, Gun Peggy S. Knudsen, Maria C. La Via, Stephanie Le Hellard, Robert D. Levitan, Dong Li, Lisa Lilenfeld, Bochao Danae Lin, Jolanta Lissowska, Jurjen Luykx, Pierre J. Magistretti, Mario Maj, Katrin Mannik, Sara Marsal, Christian R. Marshall, Morten Mattingsdal, Sara McDevitt, Peter McGuffin, Andres Metspalu, Ingrid Meulenbelt, Nadia Micali, Karen Mitchell, Alessio Maria Monteleone, Palmiero Monteleone, Benedetta Nacmias, Marie Navratilova, Ioanna Ntalla, Julie K. O’Toole, Roel A. Ophoff, Leonid Padyukov, Aarno Palotie, Jacques Pantel, Hana Papezova, Dalila Pinto, Raquel Rabionet, Anu Raevuori, Nicolas Ramoz, Ted Reichborn-Kjennerud, Valdo Ricca, Samuli Ripatti, Franziska Ritschel, Marion Roberts, Alessandro Rotondo, Dan Rujescu, Filip Rybakowski, Paolo Santonastaso, André Scherag, Stephen W. Scherer, Ulrike Schmidt, Nicholas J. Schork, Alexandra Schosser, Jochen Seitz, Lenka Slachtova, P. Eline Slagboom, Margarita C.T. Slof-Op’t Landt, Agnieszka Slopien, Sandro Sorbi, Beata Świątkowska, Jin P. Szatkiewicz, Ioanna Tachmazidou, Elena Tenconi, Alfonso Tortorella, Federica Tozzi, Janet Treasure, Artemis Tsitsika, Marta Tyszkiewicz-Nwafor, Konstantinos Tziouvas, Annemarie A. van Elburg, Eric F. van Furth, Gudrun Wagner, Esther Walton, Elisabeth Widen, Eleftheria Zeggini, Stephanie Zerwas, Stephan Zipfel, Andrew W. Bergen, Joseph M. Boden, Harry Brandt, Steven Crawford, Katherine A. Halmi, L. John Horwood, Craig Johnson, Allan S. Kaplan, Walter H. Kaye, James Mitchell, Catherine M. Olsen, John F. Pearson, Nancy L. Pedersen, Michael Strober, Thomas Werge, David C. Whiteman, D. Blake Woodside, Jakob Grove, Anjali K. Henders, Janne T. Larsen, Richard Parker, Liselotte V. Petersen, Jennifer Jordan, Martin A. Kennedy, Andreas Birgegård, Paul Lichtenstein, Claes Norring, Mikael Landén, Preben Bo Mortensen, Renato Polimanti, Jeanette N. McClintick, Amy E. Adkins, Fazil Aliev, Silviu-Alin Bacanu, Anthony Batzler, Sarah Bertelsen, Joanna M. Biernacka, Tim B. Bigdeli, Li-Shiun Chen, Toni-Kim Clarke, Franziska Degenhardt, Anna R. Docherty, Alexis C. Edwards, Jerome C. Foo, Louis Fox, Josef Frank, Laura M. Hack, Annette M. Hartmann, Sarah M. Hartz, Stefanie Heilmann-Heimbach, Colin Hodgkinson, Per Hoffmann, Jouke-Jan Hottenga, Bettina Konte, Jari Lahti, Marius Lahti-Pulkkinen, Dongbing Lai, Lannie Ligthart, Anu Loukola, Brion S. Maher, Hamdi Mbarek, Andrew M. McIntosh, Matthew B. McQueen, Jacquelyn L. Meyers, Yuri Milaneschi, Teemu Palviainen, Roseann E. Peterson, Euijung Ryu, Nancy L. Saccone, Jessica E. Salvatore, Sandra Sanchez-Roige, Melanie Schwandt, Richard Sherva, Fabian Streit, Jana Strohmaier, Nathaniel Thomas, Jen-Chyong Wang, Bradley T. Webb, Robbee Wedow, Leah Wetherill, Amanda G. Wills, Hang Zhou, Jason D. Boardman, Danfeng Chen, Doo-Sup Choi, William E. Copeland, Robert C. Culverhouse, Norbert Dahmen, Louisa Degenhardt, Benjamin W. Domingue, Mark A. Frye, Wolfgang Gäbel, Caroline Hayward, Marcus Ising, Margaret Keyes, Falk Kiefer, Gabrielle Koller, John Kramer, Samuel Kuperman, Susanne Lucae, Michael T. Lynskey, Wolfgang Maier, Karl Mann, Satu Männistö, Bertram Müller-Myhsok, Alison D. Murray, John I. Nurnberger, Ulrich Preuss, Katri Räikkönen, Maureen D. Reynolds, Monika Ridinger, Norbert Scherbaum, Marc A. Schuckit, Michael Soyka, Jens Treutlein, Stephanie H. Witt, Norbert Wodarz, Peter Zill, Daniel E. Adkins, Dorret I. Boomsma, Laura J. Bierut, Sandra A. Brown, Kathleen K. Bucholz, E. Jane Costello, Harriet de Wit, Nancy Diazgranados, Johan G. Eriksson, Lindsay A. Farrer, Tatiana M. Foroud, Nathan A. Gillespie, Alison M. Goate, David Goldman, Richard A. Grucza, Dana B. Hancock, Kathleen Mullan Harris, Victor Hesselbrock, John K. Hewitt, Christian J. Hopfer, William G. Iacono, Eric O. Johnson, Victor M. Karpyak, Kenneth S. Kendler, Henry R. Kranzler, Kenneth Krauter, Penelope A. Lind, Matt McGue, James MacKillop, Pamela A.F. Madden, Hermine H. Maes, Patrik K.E. Magnusson, Elliot C. Nelson, Markus M. Nöthen, Abraham A. Palmer, Brenda W.J.H. Penninx, Bernice Porjesz, John P. Rice, Marcella Rietschel, Brien P. Riley, Richard J. Rose, Pei-Hong Shen, Judy Silberg, Michael C. Stallings, Ralph E. Tarter, Michael M. Vanyukov, Scott Vrieze, Tamara L. Wall, John B. Whitfield, Hongyu Zhao, Benjamin M. Neale, Tracey D. Wade, Andrew C. Heath, Grant W. Montgomery, Nicholas G. Martin, Patrick F. Sullivan, Jaakko Kaprio, Gerome Breen, Joel Gelernter, Howard J. Edenberg, Cynthia M. Bulik, Arpana Agrawal

**Author notes:** Joint last authors (Cynthia M. Bulik, Arpana Agrawal). **Correspondence**: Melissa A. Munn-Chernoff, PhD, Department of Psychiatry, University of North Carolina at Chapel Hill 101 Manning Drive, Campus Box 7160 Chapel Hill, NC 27599.

## Abstract

Eating disorders and substance use disorders frequently co-occur. Twin studies reveal shared genetic variance between liabilities to eating disorders and substance use, with the strongest associations between symptoms of bulimia nervosa (BN) and problem alcohol use (genetic correlation [*r*_*g*_], twin-based=0.23-0.53). We estimated the genetic correlation between eating disorder and substance use and disorder phenotypes using data from genome-wide association studies (GWAS). Four eating disorder phenotypes (anorexia nervosa [AN], AN *with* binge-eating, AN *without* binge-eating, and a BN factor score), and eight substance-use-related phenotypes (drinks per week, alcohol use disorder [AUD], smoking initiation, current smoking, cigarettes per day, nicotine dependence, cannabis initiation, and cannabis use disorder) from eight studies were included. Significant genetic correlations were adjusted for variants associated with major depressive disorder (MDD). Total sample sizes per phenotype ranged from ~2,400 to ~537,000 individuals. We used linkage disequilibrium score regression to calculate single nucleotide polymorphism-based genetic correlations between eating disorder and substance-use-related phenotypes. Significant positive genetic associations emerged between AUD and AN (*r*_*g*_=0.18; false discovery rate *q*=0.0006), cannabis initiation and AN (*r*_*g*_=0.23; *q*<0.0001), and cannabis initiation and AN *with* binge-eating (*r*_*g*_=0.27; *q*=0.0016). Conversely, significant negative genetic correlations were observed between three non-diagnostic smoking phenotypes (smoking initiation, current smoking, and cigarettes per day) and AN *without* binge-eating (*r*_*gs*_=-0.19 to −0.23; *qs*<0.04). The genetic correlation between AUD and AN was no longer significant after co-varying for MDD loci. The patterns of association between eating disorder- and substance-use-related phenotypes highlights the potentially complex and substance-specific relationships between these behaviors.

---

A well-established phenotypic association exists between eating disorder and substance use phenotypes, with evidence for specific relations between particular types of eating disorders and substance use disorders. The prevalence of an alcohol use disorder (AUD) is greater among individuals with bulimia nervosa (BN) and binge-eating disorder (BED) than individuals with anorexia nervosa (AN) or healthy controls (Gadalla and Piran, 2007, Root *et al.*, 2010). Similarly, individuals with BN or BED are at increased risk for smoking, nicotine dependence (ND) (Solmi *et al.*, 2016, Wiederman and Pryor, 1996), and cannabis use (Krug *et al.*, 2008, Wiederman and Pryor, 1996) compared with individuals with AN or healthy controls, though these results are not consistent (Root *et al.*, 2010). Importantly, women with the binge-eating/purging subtype of AN report a higher prevalence of AUD, smoking, ND, and cannabis use than women with the restricting subtype of AN (Anzengruber *et al.*, 2006, Krug *et al.*, 2008, Root *et al.*, 2010). Thus, binge eating—a transdiagnostic symptom defined as eating a large amount of food in a short period of time while experiencing loss of control—may be a key component of the observed association.

However, prior research has only partially addressed whether binge eating is the critical eating disorder symptom in the comorbidity, especially across different milestones of substance use (i.e., initiation through substance use disorder) and across a variety of substances (i.e., alcohol, nicotine, and cannabis). It is crucial to elucidate shared sources for these associations because of the increased morbidity and mortality associated with comorbid presentations (Duncan *et al.*, 2006, Franko *et al.*, 2013) and because improvements in one disorder may exacerbate (or weaken) symptoms of the other disorder (Center on Addiction and Substance Abuse, 2003). Refining our understanding of these associations could improve prevention and treatment approaches for these debilitating disorders, their comorbidity, and their sequelae.

Accumulating findings from twin studies implicate shared genetic factors between eating disorder- and substance-use-related phenotypes. The strongest reported association is between BN symptoms, including binge eating, and problem alcohol use (Munn-Chernoff and Baker, 2016), with a genetic correlation (*r*_*g*_) ranging from 0.23 to 0.53 (Baker *et al.*, 2010, Baker *et al.*, 2017, Munn-Chernoff *et al.*, 2013, Munn-Chernoff *et al.*, 2015, Slane *et al.*, 2012, Trace *et al.*, 2013). Although there has been less focus on the genetic associations between BN symptoms and regular smoking and BN symptoms and illicit drug use disorder, twin-based *r*_*g*_s of 0.35 and approximately 0.38, respectively, have been reported (Baker *et al.*, 2007, Baker *et al.*, 2010). A paucity of information exists regarding whether less problematic aspects of substance use exhibit a significant *r*_*g*_ with eating disorder phenotypes. The impact of genetic factors influencing this comorbidity may significantly increase once an individual has progressed to problematic alcohol use, as genetic effects are more prominent in problem substance use, such as abuse and dependence, than with the initiation and general use of substances (Heath *et al.*, 1997, Rhee *et al.*, 2003, True *et al.*, 1997, van den Bree *et al.*, 1998). No study has comprehensively examined a range of eating disorder- and substance-use-related phenotypes to determine whether the *r*_*g*_ varies with different aspects of substance use and whether the *r*_*g*_ varies depending on the eating disorder and substance examined.

Recent advances in genomic methods allow for an assessment of *r*_*g*_ using existing genome-wide association study (GWAS) summary statistics. Unlike twin studies, these genome-wide methods allow for use of unrelated cases and controls, which typically yield large sample sizes (i.e., in the tens to hundreds of thousands). One such method is linkage disequilibrium score regression (LDSC; Bulik-Sullivan *et al.*, 2015a, Bulik-Sullivan *et al.*, 2015b), which estimates single-nucleotide polymorphism (SNP)-based heritability and *r*_*g*_ between phenotypes. Of particular relevance to low prevalence phenotypes, such as AN, estimation of SNP-based *r*_*g*_ does not require both phenotypes to be measured in the same individual, meaning that independent studies assessing only one phenotype can be jointly examined.

Thus, the aim of the current study is to estimate SNP-based *r*_*g*_s between eating disorder- and substance-use-related phenotypes based upon summary statistics from the largest published eating disorder GWAS and existing GWAS encompassing a range of substance-use-related phenotypes. This study examines shared SNP-based genetic risk between eating disorder and multiple substance-use-related phenotypes (i.e., alcohol, nicotine, and cannabis), using robust data from twin studies to shape our expectations. First, we hypothesize that the strongest SNP-based *r*_*g*_ will be between eating disorder phenotypes that have binge eating as a core symptom and alcohol use phenotypes (Munn-Chernoff and Baker, 2016). Second, we hypothesize that for binge-eating-related phenotypes, the SNP-based *r*_*g*_ will be lowest when examining typical alcohol consumption and highest when assessing AUD (Munn-Chernoff and Baker, 2016). Because we have less information from twin studies about genetic associations between liabilities to eating disorders and tobacco (nicotine) and cannabis use-related phenotypes, we do not forward any hypotheses for these substances. Finally, prior studies document robust genetic associations between major depressive disorder (MDD) and both eating disorders and substance-use-related phenotypes (e.g., Kranzler *et al.*, 2019; Liu *et al.*, 2019; Watson *et al.*, 2019). We hypothesized that *r*_*g*_s between eating disorders and substance use and disorder would be attenuated when accounting for variants associated with MDD. Findings from this study could yield important information about this clinically challenging pattern of comorbidity (Gregorowski *et al.*, 2013), ultimately suggesting biologically informed prevention efforts and improved treatments for patients presenting with these dual diagnoses.

## Method

### Participants

We included summary statistics from two existing GWAS of eating disorder phenotypes where particiants were primarily of European ancestry (Wade *et al.*, 2013, Watson *et al.*, 2019) and using data from individuals of European ancestry from six existing GWAS of substance-use-related phenotypes (Demontis *et al.*, 2019, Hancock *et al.*, 2017, Kranzler *et al.*, 2019, Liu *et al.*, 2019, Pasman *et al.*, 2018, Walters *et al.*, 2018). The eating disorder phenotypes (Table 1) included a diagnosis of AN (which was further parsed into AN *with* binge-eating or AN *without* binge-eating), and a BN factor score derived from the Eating Disorder Examination (EDE; Fairburn and Cooper, 1993). The EDE is a well-established, structured clinical interview used to determine eating disorder diagnoses. We did not examine BN or BED diagnoses because there are currently no published GWAS for either disorder; thus, the EDE-BN factor score represents the closest to a GWAS of BN available. Substance-use-related phenotypes ranged from typical use (e.g., drinks per week, smoking initiation, and cannabis initiation) to substance use disorder (i.e., AUD, ND, and cannabis use disorder [CUD]). Sample sizes for the phenotypes ranged from 2,442 (BN factor) to 537,349 (drinks per week) individuals. Table 2 provides individual study details.

**Table 1.** Eating disorder-related phenotype descriptions.

| Phenotype | Definitions |
| --- | --- |
| Anorexia nervosa (AN)<sup>a</sup> | Diagnostic criteria included:<br>1. BMI less than minimally expected<br>2. Intense fear of gaining weight<br>3. Weight or shape disturbance, undue influence of weight or shape, or denial of the seriousness of the disorder |
| AN *with* binge-eating<sup>b</sup> | Individuals with AN who also engaged in binge eating episodes, defined as eating a large amount of food in a short period of time while having a sense of loss of control over the eating episode. The binge eating episodes must have occurred at least twice a week for three months. |
| AN *without* binge-eating<sup>b</sup> | Individuals with AN who did not engage in binge eating episodes. |
| Bulimia nervosa (BN)<sup>c</sup> factor | Derived from a factor analysis that included the following items:<br>1. Reporting self-induced vomiting to control body weight<br>2. Reporting suffering from or being treated for binge eating<br>3. Reporting suffering from or being treated for bulimia |

**Note:** <sup>a</sup>A fourth diagnostic criterion for AN includes amenorrhea. However, amenorrhea was excluded as a required criterion for cases in the Psychiatric Genomics Consortium datasets since it is no longer a diagnostic criterion in the DSM-5. <sup>b</sup>The DSM and ICD include two subtypes of anorexia nervosa (AN)—a binge-eating/purging subtype and a restricting subtype. Although it would have been ideal to examine differences between the AN binge-eating/purging subtype and AN restricting subtype, this was not possible with current Psychiatric Genomics Consortium data. However, there was sufficient information about presence or absence of binge eating, which resulted in creating the AN *with* binge-eating and AN *without* binge-eating subtypes. <sup>c</sup>Bulimia nervosa is defined as: 1) recurrent episodes of binge eating; 2) recurrent inappropriate compensatory behaviors (e.g., self-induced vomiting, laxative use) to prevent weight gain; 3) the binge eating and inappropriate compensatory behaviors occurring an average of twice a week for three months; 4) having undue influence of body weight and shape; and 5) disturbance not occurring during AN.

**Table 2.**
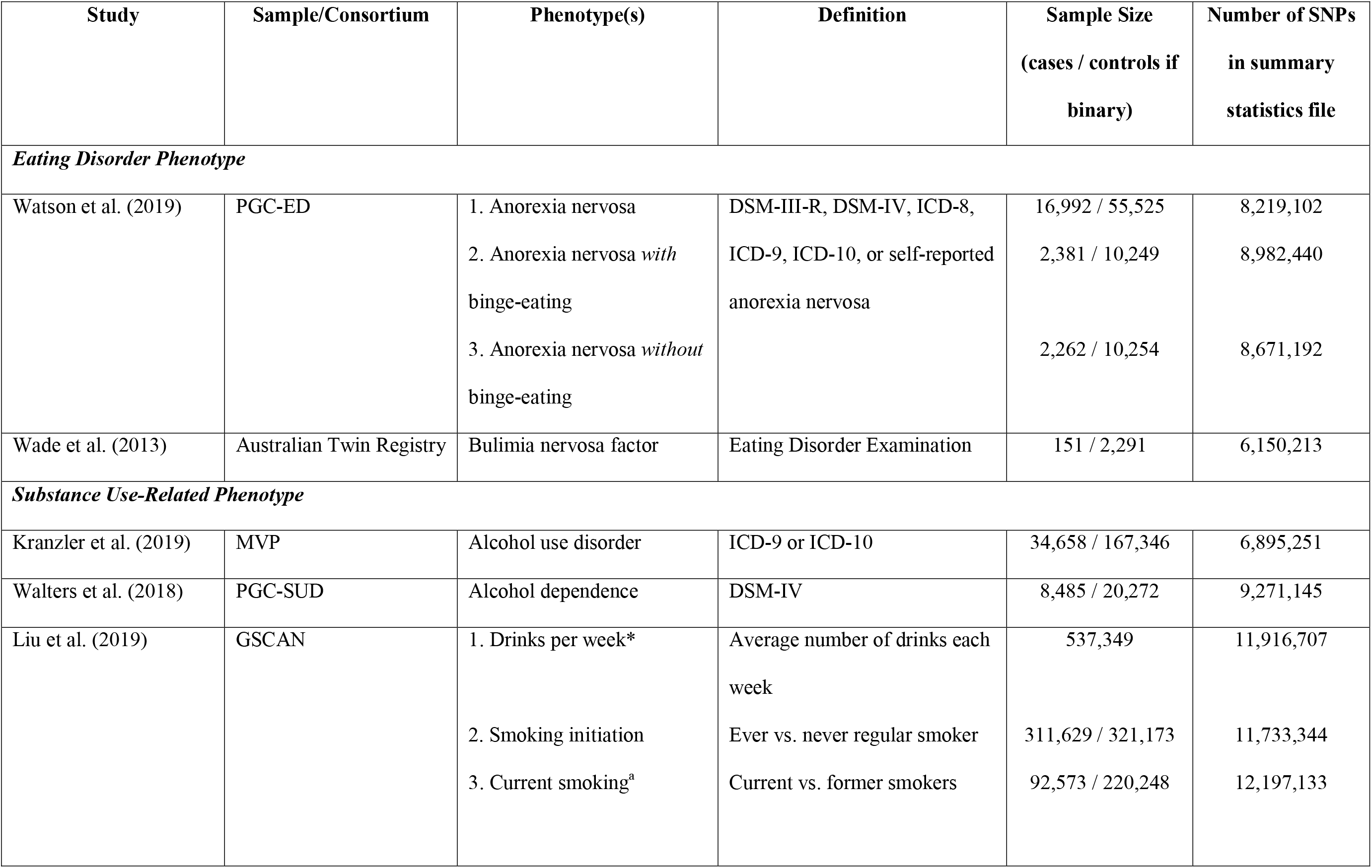
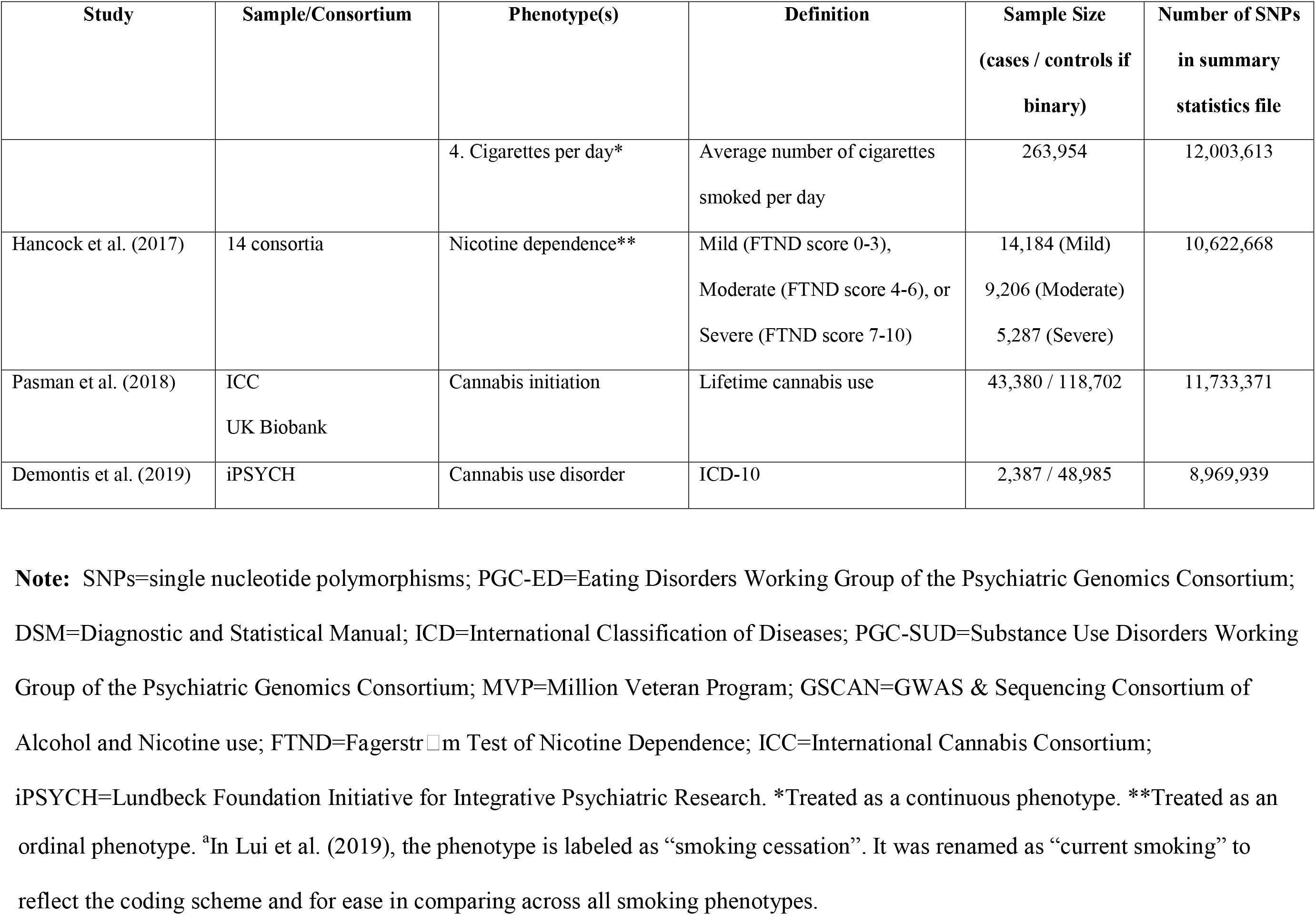
Details of samples included in analyses.

### Statistical Analysis

We used LDSC (Bulik-Sullivan *et al.*, 2015a, Bulik-Sullivan *et al.*, 2015b) to evaluate SNP-based genetic correlations (*r*_*g*_) between samples. This method uses the linkage disequilibrium (LD) structure of the genome to estimate the distribution of effect sizes for individual SNPs as a function of their LD score. Under a polygenic model, causal SNPs are likely to be overrepresented in higher LD score bins (i.e., including additional SNPs in high LD) such that associations with SNPs in these LD bins will make stronger contributions to the phenotypic variation under study. This polygenic distribution of effect sizes across LD score bins provides an estimate of SNP-based heritability, i.e., the proportion of phenotypic variance that is attributable to the aggregate effects of genome-wide SNPs. The correlation of effect sizes across LD bins between two phenotypes then provides an estimate of SNP-based *r*_*g*_.

Genetic correlations range from −1 to +1, where the sign indicates that the same genetic factors are contributing to variation in the target traits in *opposite* or *same* directions, respectively. The LDSC intercept for the genetic covariance provides evidence about sample overlap across two traits. SNPs (MAF>0.01) found in the HapMap3 EUR population were used to calculate LD scores. We used the false discovery rate (FDR; Benjamini and Hochberg, 1995) to correct for multiple testing (*n*=66 tests; *q*<0.05). Finally, post-hoc analyses examined whether significant differences between two *r*_*g*_s existed, using the jackknife procedure implemented through LDSC (Bulik-Sullivan *et al.*, 2015b).

For significant *r*_*g*_s detected in LDSC, multi-trait-based conditional and joint analysis using GWAS summary data (mtCOJO; Zhu *et al.*, 2018) was used to condition both input GWAS (e.g., AN and AUD) for variants associated with MDD at *p*<5×10^−7^ (Wray *et al.*, 2018). LDSC was used to compute *r*_*g*_s using the resulting genome-wide summary statistics for each trait after adjustment for MDD variants to examine whether conditioning on MDD would affect the observed genetic relationships. Further, for significant *r*_*g*_s detected in LDSC where both the individual eating disorder- and substance-use-related phenotypes each had ~10 or more significant GWAS SNPs (Zhu *et al.*, 2018), bidirectional Mendelian randomization (MR) analyses (Smith and Ebrahim, 2003) were conducted using Generalised Summary-data-based Mendelian Randomisation (GSMR; Zhu *et al.*, 2018) to preliminarily investigate potentially causal relationships between liabilities to these phenotypes. As GSMR requires a reference sample with individual genotypes to account for LD, we used the 1000 Genomes Phase 3 European ancestries sample as our reference panel (1000 Genomes Project Consortium *et al.*, 2015). SNPs with evidence of horizontal pleiotropy were excluded (using the HEIDI-Outlier method; default *p*-value threshold=0.01). We only included genome-wide significant SNPs (*p*<5×10^−8^) as possible instruments, and SNPs were clumped to ensure independence (i.e., amongst a set of genome-wide significant SNPs correlated at *r*^*2*^>0.05 within a 1-Mb window, only the SNP with the lowest *p*-value was retained). As MR analyses are sensitive to sample overlap, we scanned the published papers to determine which samples were included in the discovery GWAS, and any known samples in common across the two discovery GWAS were excluded from the eating disorder GWAS and summary statistics were regenerated for MR analyses.

## Results

The overall SNP-based heritability for the eating disorder phenotypes ranged from 0.20 to 0.39, whereas the corresponding heritabilities for the substance-use-related phenotypes ranged from 0.03 to 0.35 (**Supplemental Table 1**). Figure 1 and **Supplemental Table 1** show the genetic correlations (*r*_*g*_s) between all four eating disorder phenotypes and eight substance-use-related phenotypes. Broadly speaking, there were significant *r*_*g*_s across substance-use-related phenotypes, ranging from 0.21 (AUD and cigarettes per day) to 0.70 (drinks per week and AUD). Cannabis initiation risk was not significantly genetically correlated with cigarettes per day or ND. For the remaining results, we focus on previously unexplored associations of interest in this study—correlations between eating disorder- and substance-use-related phenotypes. For these associations, the genetic covariance intercepts ranged from −0.03 (standard error [SE]=0.01; AN and cannabis initiation) to 0.01 (SE=0.01; AN and CUD), indicating some sample overlap (or low-level confounding) existed (Yengo *et al.*, 2018), although the LDSC approach parses this overlap from the *r*_*g*_ estimation.

**Figure 1.**
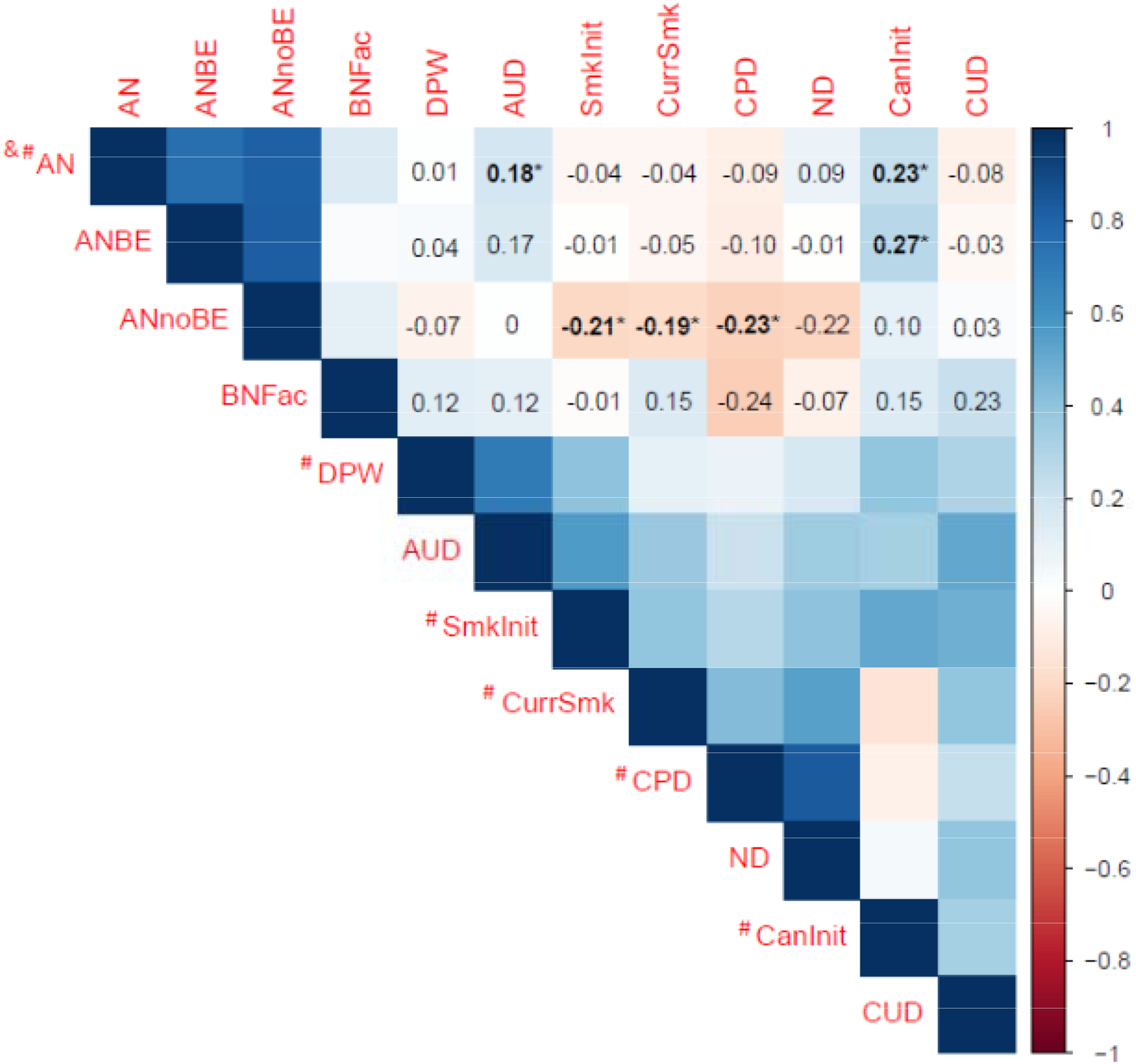
Genetic correlations between eating disorder subtypes and substance-use-related phenotypes. AN=anorexia nervosa; ANBE=anorexia nervosa *with* binge-eating; ANnoBE=anorexia nervosa *without* binge-eating; BNFac=bulimia nervosa factor score; DPW=drinks per week; AUD=alcohol use disorder; SmkInit=smoking initiation; CurrSmk=current smoking; CPD=cigarettes per day; ND=nicotine dependence; CanInit=cannabis initiation; CUD=cannabis use disorder. # indicates known or potential sample overlap with UK Biobank; & indicates known sample overlap with iPSYCH. Bolded and * values denote significant genetic correlations after correcting for multiple comparisons using False Discovery Rate (*n* tests=66; *q*<0.05).

Significant positive *r*_*g*_s were observed for alcohol- and cannabis-use-related phenotypes. First, the *r*_*g*_ was significant between AN and AUD (*r*_*g*_=0.18; SE=0.05; *q*=0.0006), but not between AN and drinks per week (*r*_*g*_=0.01; SE=0.03; *q*=0.91), suggesting that genetic factors that increase risk for AN also increase risk for AUD, but little evidence exists for shared genetic risk between AN and typical alcohol consumption. These two correlations significantly differed from each other (z-score=3.51, *p*=0.0005). Intriguingly, there was a significant difference in *r*_*g*_s for AN and AUD versus AN *without* binge-eating and AUD (z-score=2.28, *p*=0.02), but not for AN and AUD versus AN *with* binge-eating and AUD (z-score=0.23, *p*=0.82). No significant association between the BN factor, which included items pertaining to both binge eating and compensatory behaviors, and either alcohol-use-related phenotype was observed.

Second, the significant *r*_*g*_ between AN and cannabis initiation was 0.23 (SE=0.04, *q*<0.0001), and the significant *r*_*g*_ between AN *with* binge-eating and cannabis initiation was 0.27 (SE=0.08, *q*=0.0017), indicating that genetic factors that increase the risk for AN also increase the risk for cannabis initiation. However, cannabis initiation was not significantly correlated with the BN factor (*r*_*g*_=0.15, SE=0.18, *q*=0.57) or with AN *without* binge-eating (*r*_*g*_=0.10, SE=0.08, *q*=0.31). No significant associations were observed between any eating disorder phenotype and CUD (*r*_*g*_s=-0.08-0.23; SEs=0.01; *q*s≤0.57). Post-hoc analyses revealed significant differences in the *r*_*g*_s for AN and cannabis initiation versus AN and CUD (z-score=2.70, *p*=0.01). However, the *r*_*g*_ between AN *with* binge-eating and cannabis initiation, while significant, was statistically different from the *r*_*g*_ between AN *with* binge-eating and CUD.

Conversely, for smoking phenotypes, significant correlations were only observed for the AN *without* binge-eating subtype. Smoking initiation (*r*_*g*_=-0.21, SE=0.06, *q*=0.0006), current smoking (referred to as smoking cessation in Liu *et al.*, 2019)^1^ (*r*_*g*_=-0.19, SE=0.08, *q*=0.03), and cigarettes per day (*r*_*g*_=-0.23, SE=0.07, *q*=0.003) were significantly and negatively associated with AN *without* binge-eating. Although the correlation between ND and AN *without* binge-eating was in the same direction as the other smoking phenotypes, it was not significant (*r*_*g*_=- 0.22, SE=0.12, *q*=0.14). The *r*_*g*_s for AN diagnosis and each of the three non-diagnostic smoking traits versus AN *without* binge-eating and these same smoking traits all differed significantly from each other (z-scores ranged from −3.22 to −2.11; *p*-values≤0.04).

After conditioning the AN and AUD GWAS summary statistics for the MDD GWAS summary statistics, the positive *r*_*g*_ between AN and AUD was attenuated (*r*_*g*_=0.07; SE=0.05, *q*=0.125; **Supplemental Table 2**) and significantly lower than the unadjusted *r*_*g*_ (z-score=2.48, *p*=0.01). In contrast, after conditioning the AN *with* binge-eating and cannabis initiation GWAS for MDD, the resulting *r*_*g*_ was marginally smaller but remained significant after correction for multiple tests (*r*_*g*_=0.21, SE=0.08, *q*=0.016). After conditioning for the MDD GWAS, *r*_*g*_s between AN *without* binge-eating and smoking initiation, current smoking, and cigarettes per day remained significant and modestly increased in magnitude (*r*_*g*_s=-0.27 to −0.31; SEs=0.05 to 0.09; *q*s<0.0009).

Because the AN GWAS and cannabis initiation GWAS each identified eight significant loci (Pasman *et al.*, 2018, Watson *et al.*, 2019), and the two studies comprising the AUD sample identified 10 significant loci (Kranzler *et al.*, 2019, Walters *et al.*, 2018), we also conducted exploratory follow-up MR analyses for AN-AUD and AN-cannabis initiation to examine whether there might be evidence of a causal relationship, given their significant *r*_*g*_. We used summary statistics from a subset of the AN GWAS that did not include the UK Biobank cohort for the AN-cannabis initiation analysis, as this cohort overlapped with the cannabis initiation sample (AN subset without the UK Biobank cohort, *N*_*cases*_=16,224, *N*_*controls*_=52,460). We did not obtain an estimate for the cannabis inititiation to AN direction of effect because only four SNPs were available after clumping, all of which were excluded for potential pleiotropy by the HEIDI-outlier analysis. There was no evidence of a causal relationship for any comparison (*p*>0.05; see **Supplemental Table 3**).

## Discussion

Using existing GWAS data, we investigated genetic associations between liabilities to four eating disorder and eight substance-use-related phenotypes spanning initiation and typical use to substance use disorder. We found differential patterns of association between the two diagnostic categories for eating disorders, suggesting that differences between AN subtypes with substance-use-related phenotypes may point toward substance-specific genetic relationships. Additionally, there may be some degree of symptom overlap contributing to these associations.

Three main patterns emerged. First, in line with prior twin studies, we observed a positive genetic correlation (*r*_*g*_) between problem alcohol use (i.e., AUD) and AN diagnosis. Second, we estimated positive *r*_*g*_s between cannabis initiation and AN diagnosis, as well as cannabis initiation and the AN *with* binge-eating subtype. This is a novel finding not previously examined in twin research. The positive genetic associations suggest that some genetic loci are influencing these traits in the same direction. Second, negative *r*_*g*_s emerged between the three smoking phenotypes from the GWAS & Sequencing Consortium of Alcohol and Nicotine use (GSCAN) cohort and AN *without* binge-eating, but not with the other three eating disorder phenotypes. These negative *r*_*g*_s indicate that some of the loci influencing the liability to these eating disorder and smoking phenotypes are shared, but are affecting the liability to these traits in opposite directions. Indeed, *r*_*g*_s cannot identify specific loci or underlying mechanisms that contribute to the shared risk. Nevertheless, the results provide initial evidence for differential genetic associations between the liability to varying eating disorder- and substance-use-related phenotypes.

Based on findings from twin studies, we hypothesized that: 1) the strongest SNP-based *r*_*g*_ would be between eating disorder phenotypes that have binge eating as a core symptom and alcohol use phenotypes; and 2) a significant positive *r*_*g*_ between eating disorder phenotypes with binge eating as a key symptom and AUD would emerge. In line with these hypotheses, we found a signifant genetic association between AUD and AN diagnosis, but not between typical alcohol consumption (i.e., drinks per week) and AN. No twin study has examined genetic associations between AN and alcohol-use-related phenotypes, and previous studies (Walters *et al.*, 2018, Watson *et al.*, 2019) using LDSC have not reported significant *r*_*g*_s between these traits. That we found a significant association most likely reflects the larger AN sample size in our study (from 3,495 cases and 10,982 controls to 16,992 cases and 55,525 controls), as well as combining two large existing GWAS of AUD, emphasizing the importance of increasing sample sizes for GWAS.

Importantly, the *r*_*g*_ between AN and AUD was not robust to the adjustment for MDD-associated variants. MDD is amongst the most prominent comorbidities in individuals with AN and AUD (American Psychological Association, 2013), and GWAS for both traits document strong *r*_*g*_s between MDD and these disorders (Kranzler *et al.*, 2019; Walters *et al.*, 2019; Watson *et al.*, 2019). Our results indicate that the three disorders share genetic underpinnings. We cannot discount the possibility of a genetic relationship between AN and AUD that is distinct from MDD; however, much larger sample sizes may be required to detect such an association.

Intriguingly, although we did not detect a significant *r*_*g*_ for AN *with* binge-eating with AUD, the point estimate for the *r*_*g*_ between AUD and AN *with* binge eating was similar to that for AUD and AN diagnosis (0.17 vs. 0.18, respectively) and higher than AUD and AN *without* BE (0.01). Sample sizes for these AN subtypes were smaller than for AN diagnosis; however, the two subtypes included approximately equal numbers of cases and controls. Indeed, binge eating was assessed in such a way that we were unable to tease apart purging behaviors, and AN diagosis is heterogenous even within subytpes. Therefore, binge eating may be one plausible key component of the observed genetic association. For example, binge eating has been shown to activate brain reward circuitry in a similar manner to substances (Kaye *et al.*, 2013, Volkow *et al.*, 2013), and administration of naltrexone, an opioid antagonist approved by the U.S. Food and Drug Administration for the treatment of AUD (Kranzler and Soyka, 2018), has been shown to reduce the frequency of binge-eating episodes among individuals with an eating disorder (Jonas and Gold, 1988, Stancil *et al.*, in press). We did not detect a significant *r*_*g*_ with the BN factor score as well, although that GWAS was relatively underpowered. Thus, our findings highlight the importance of expanding GWAS to include BN and BED, where a core symptom of both disorders is binge eating, to elucidate whether binge eating is a critical eating disorder symptom in the comorbidity with AUD and to home in on the relevant shared mechanisms.

The significant genetic associations between cannabis initiation and AN are novel, yet consistent with the negative genetic association between cannabis use and BMI, and with observational (Pasman *et al.*, 2018) and experimental (Di Marzo and Matias, 2005, Volkow *et al.*, 2017) studies regarding the role of endocannabinoids in appetite regulation, energy expenditure, stress, and reward. One of the principal psychoactive agents of cannabis, delta-9-tetrahydrocannabinol (THC), a partial agonist of the endogenous cannabinoid 1 (CB1) receptor, is presumed to be orexigenic and may acutely increase appetite and food intake, contributing to its potential role as an appetite stimulant in patients with an anorexia or cachexia syndrome (Reuter and Martin, 2016) due to a disease (e.g., HIV AIDS) or in response to treatment (e.g., chemotherapy). An antagonist of the CB1 receptor was previously tested as a highly promising anti-obesity medication (Rimonabant, SR141716). Further, the endocannabinoid anandamide has been shown to be elevated in individuals with acute AN (Monteleone and Maj, 2013), indicating disruption in food-related reward and eating behavior regulation. Animal and human studies have also provided initial evidence for the therapeutic effectiveness of cannabinoid agonists in treating eating disorders (Andries *et al.*, 2014, Avraham *et al.*, 2017). It is also likely that individuals with high genetic liability to AN are less likely to experiment with a substance that has a documented hyperphagia component. Thus, there is evidence of a complex biological relationship between cannabis use and eating disorders, as well as BMI. Nonetheless, the preliminary MR analyses did not provide persuasive support for causal relationships, in either direction, between genetic liability to cannabis initiation and AN. The small sample sizes and small number of SNPs used as instruments, due to the low number of independent genome-wide significant loci and exclusion of overlapping SNPs that were not ambiguous and palindromic, may have significantly limited power to make causal inferences.

Finally, the significant negative *r*_*g*_s between three tobacco-smoking phenotypes— smoking initiation, current smoking, and cigarettes per day—and AN *without* binge-eating are intriguing, suggesting that AN *without* binge-eating and tobacco-smoking behaviors are alternate expressions of shared mechanisms. Phenotypic studies are inconsistent about the association between the restricting subtype of AN and smoking. Some studies suggest that individuals with restricting AN have a higher prevalence of various smoking phenotypes than controls (Krug *et al.*, 2008), whereas other studies indicate no significant difference between the two groups (Anzengruber *et al.*, 2006). A recent meta-analysis did not find differences in the odds of lifetime smoking between individuals with AN and healthy controls (Solmi *et al.*, 2016), yet the authors did not assess differences by AN subtype. Individuals with AN may smoke as a way to control or lose weight (White, 2011), and temporary weight gain does occur with smoking cessation (Filozof *et al.*, 2004). However, a positive phenotypic correlation need not be accompanied by a *r*_*g*_ in the same direction (or genetic contributors to the phenotypic association at all). Still, there is plausible support for the negative *r*_*g*_. Although not significant, a negative *r*_*g*_ between smoking and AN has been reported (Bulik-Sullivan *et al.*, 2015a, Watson *et al.*, 2019). Notably, our study includes individuals from these earlier reports and extends findings by including larger sample sizes for both AN and smoking phenotypes. Unfortunately, there are no twin studies of AN or AN-like traits and smoking with which to compare findings.

One explanation for the negative genetic association is that it is due to a third, underlying variable influencing both AN *without* binge-eating and smoking. We tested for the potential role of variants associated with MDD and found the *r*_*g*_s to be robust to that adjustment. In the largest GWAS of smoking phenotypes, positive *r*_*g*_s were also observed between smoking initiation and cigarettes per day with multiple cardiometabolic traits, including type 2 diabetes and fasting glucose (Liu *et al.*, 2019). These same metabolic traits were negatively genetically correlated with AN (Duncan *et al.*, 2017, Watson *et al.*, 2019). Thus, the patterns of *r*_*g*_s might point to metabolic, rather than psychiatric, factors in influencing the apparent genetic association between smoking phenotypes and AN. However, the associations could also reflect adoption of unhealthy lifestyles that promote obesity and are correlated with smoking. In addition, the *r*_*g*_s between smoking and BMI, as well as AN and BMI, could reflect underlying disinhibitory pathways, as variants associated with BMI show enrichment in the central nervous system (Goodarzi, 2018). The current approach is not designed to disentangle these putative etiological mechanisms, but our findings do encourage careful study of the specific relationships between eating and substance use disorders.

Substance use and substance use disorders are partially distinct, and although excessive substance use is a necessary component of substance use disorders, the latter is associated with psychological and physiological impairment related to excess use and aspects of loss of control over the behavior. Consistent with our findings for alcohol, accumulating evidence suggests that genetic liability to other psychiatric traits (e.g., schizophrenia) is strongly correlated with liability to substance use disorders (e.g., AUD) but not substance use (e.g., alcohol consumption; Kranzler *et al*., 2019; Liu *et al.*, 2019; Walters *et al*., 2018). Genetic liability to alcohol use has also been correlated with liabilities to psychiatric disorders (e.g., MDD) in opposite directions depending on level of involvement (Kranzler *et al.*, 2019). However, we did not find similar elevations in *r*_*g*_s when contrasting ever smoking and ND, nor comparing cannabis initiation to CUD. It is possible that the lack of genetic overlap between AN and ND, as well as AN and CUD, is related to the relatively modest sample size of those discovery GWAS. A similar non-significant *r*_*g*_ was noted for AUD when the Walters *et al.* (2018) alcohol dependence GWAS was used as the sole source of summary statistics for problem drinking in the current study. Several other explanations for this divergence in findings exist. For instance, for tobacco, the highly addictive nature of nicotine may result in convergence in genomic effects on earlier and later stages of smoking (i.e., a much larger proportion of those who ever smoke become dependent compared with the proportion of those who drink alcohol and develop AUD). For cannabis, given its lower addictive potential, we might have expected stronger associations with CUD than with cannabis initiation. In addition to the considerably smaller sample size of the CUD GWAS, the association with cannabis initiation could also be attributed to the small number of cohorts in that discovery GWAS that included individuals with a high likelihood of CUD. It is also possible that the relationship between AN and cannabis use is distinct and that earlier, but not later stages of cannabis use are genetically related to liability to AN. Future studies should consider the multi-stage nature of substance use and misuse when examining cross-trait correlations.

This is the largest and most comprehensive assessment of shared genetic risk between eating disorder- and substance-use-related phenotypes to date, using existing GWAS data from large cohorts (up to ~537,000 individuals per phenotype). We were able to separately assess approximate AN subtypes (i.e., *with* binge-eating vs. *without* binge-eating) to evaluate the extent to which binge eating, in the context of AN, may share genetic risk with substance-use-related phenotypes. Using these large datasets—many of which are publicly available—allows for the rapid development of scientific knowledge regarding the underlying etiology of psychiatric disorder and substance use comorbidity. Nevertheless, some limitations exist. First, sample sizes for the BN factor score and CUD GWAS were relatively small compared with the other GWAS, resulting in large standard errors and low power. Second, we were unable to uniformally examine sex differences in these *r*_*g*_s. Since the prevalence of eating disorders is higher in women than men, and the prevalence of substance use disorders is higher in men than women (American Psychiatric Association, 2013), it will be important to explore possible sex differences in genetic associations as the GWAS data become available. Notably, we previously did not find evidence for sex differences in the *r*_*g*_ between binge eating and problem alcohol use (Munn-Chernoff *et al.*, 2013). Finally, SNP coverage was limited in the earlier GWAS of the BN factor score because that study used older genotyping platforms and imputation panels that included fewer SNPs than current imputation panels. The Eating Disorders and Substance Use Disorders Working Groups of the Psychiatric Genomics Consortium (PGC) are continuously adding samples and releasing data freezes with incrementally larger sample sizes, while collecting information on multiple substances (e.g., opioids). Thus, in coming years, the statistical power is expected to increase for AN (including the *with* and *without* binge-eating subtypes), BN, and BED, as well as AUD, ND, and CUD, from within and outside the PGC. This will allow for a more refined assessment of specific eating disorder symptoms, including binge eating, in relation to substance-use-related phenotypes.

In conclusion, findings from this study suggest that the shared sources of variation in liabilities to eating disorder and substance-use-related phenotypes are not consistent across traits or levels of substance involvement, extending results from twin studies to a genome-wide SNP approach. Despite the typically high co-occurrence of alcohol, tobacco, and cannabis use, and their genetic overlap (Pasman *et al.*, 2018), the differential patterns seen between the eating disorder- and substance-use-related phenotypes highlight the uniqueness and complexity of their shared etiology. Additional research using contemporary genomic methods such as cross-disorder association studies could identify the specific loci contributing to this comorbidity. Once loci are identified, additional research that combines polygenic risk scores with measured environmental constructs could enhance the prediction, prevention, and treatment of co-occurring eating disorder- and substance-use-related traits.

## Acknowledgements

Grant support for individual authors can be found in **Supplementary Table 4**. This study included summary statistics of a genetic study on cannabis use (Pasman et al. [2018] *Nature Neuroscience*). We would like to acknowledge all participating groups of the International Cannabis Consortium, and in particular, the members of the working group including Joelle Pasman, Karin Verweij, Nathan Gillespie, Eske Derks, and Jacqueline Vink. Pasman et al. (2018) included data from the UK Biobank resource under application numbers 9905, 16406, and 25331.

## Eating Disorders Working Group of the Psychiatric Genomics Consortium (PGC-ED)

We thank all study volunteers, study coordinators, and research staff who enabled this study. ANGI: The Anorexia Nervosa Genetics Initiative was an initiative of the Klarman Family Foundation. Additional support was offered by the National Institute of Mental Health. We acknowledge support from the North Carolina Translational and Clinical Sciences Institute (NC TraCS) and the Carolina Data Warehouse. PGC: We are deeply indebted to the investigators who comprise the PGC, and to the hundreds of thousands of individuals who have shared their life experiences with PGC investigators and the contributing studies. We are grateful to the Children’s Hospital of Philadelphia (CHOP), the Price Foundation Collaborative Group (PFCG), Genetic Consortium for Anorexia Nervosa (GCAN), Wellcome Trust Case-Control Consortium-3 (WTCCC-3), the Lundbeck Foundation Initiative for Integrative Psychiatric Research (iPSYCH), the QSkin Sun and Health Study, Riksät (Swedish National Quality Register for Eating Disorders), the Stockholm Center for Eating Disorders (SCÄ), LifeGene, the UK Biobank, and all PGC-ED members for their support in providing individual samples used in this study. We thank SURFsara (http://www.surf.nl) for support in using the Lisa Compute Cluster. We thank Max Lam, Institute of Mental Health, Singapore, for Ricopili consultation. This study also represents independent research partly funded by the English National Institute for Health Research (NIHR) Biomedical Research Centre at South London and Maudsley NHS Foundation Trust and King’s College London. The views expressed are those of the author(s) and not necessarily those of the NHS, the NIHR, or the English Department of Health and Social Care. High performance computing facilities were funded with capital equipment grants from the GSTT Charity (TR130505) and Maudsley Charity (980). Research reported in this publication was supported by the National Institute of Mental Health of the US National Institutes of Health under Award Number U01MH109514. The content is solely the responsibility of the authors and does not necessarily represent the official views of the U.S. National Institutes of Health.

## Substance Use Disorders Working Group of the Psychiatric Genomics Consortium (PGC-SUD)

The PGC-SUD receives support from the National Institute on Drug Abuse and the National Institute of Mental Health via MH109532. We gratefully acknowledge prior support from the National Institute on Alcohol Abuse and Alcoholism. Statistical analyses for the PGC were carried out on the Genetic Cluster Computer (http://www.geneticcluster.org) hosted by SURFsara and financially supported by the Netherlands Scientific Organization (NWO 480-05-003) along with a supplement from the Dutch Brain Foundation and the VU University Amsterdam. Cohort specific acknowledgements may be found in Walters et al. (2018) *Nature Neuroscience*.

## Competing Financial Interests

The authors report the following potential competing interests. O. Andreassen received a speaker’s honorarium from Lundbeck. G. Breen received grant funding and consultancy fees in preclinical genetics from Eli Lilly, consultancy fees from Otsuka, and has received honoraria from Illumina. C. Bulik served on Shire Scientific Advisory Boards; she receives author royalties from Pearson. D. Degortes served as a speaker and on advisory boards, and has received consultancy fees for participation in research from various pharmaceutical industry companies including: AstraZeneca, Boehringer, Bristol Myers Squibb, Eli Lilly, Genesis Pharma, GlaxoSmithKline, Janssen, Lundbeck, Organon, Sanofi, UniPharma, and Wyeth; he has received unrestricted grants from Lilly and AstraZeneca as director of the Sleep Research Unit of Eginition Hospital (National and Kapodistrian University of Athens, Greece). J. Hudson has received grant support from Shire and Sunovion, and has received consulting fees from DiaMentis, Shire, and Sunovion. A. Kaplan is a member of the Shire Canadian BED Advisory Board and was on the steering committee for the Shire B/educated Educational Symposium: June 15-16, 2018. J. Kennedy served as an unpaid member of the scientific advisory board of AssurexHealth Inc. M. Landén declares that, over the past 36 months, he has received lecture honoraria from Lundbeck and served as scientific consultant for EPID Research Oy. No other equity ownership, profit-sharing agreements, royalties, or patent. S. Scherer is a member of the scientific advisory board for Deep Genomics. P. Sullivan is on the Lundbeck advisory committee and is a Lundbeck grant recipient; he has served on the scientific advisory board for Pfizer, has received a consultation fee from Element Genomics, and a speaker reimbursement fee from Roche. J. Treasure has received an honorarium for participation in an EAP meeting and has received royalties from several books from Routledge, Wiley, and Oxford University press. T. Werge has acted as a lecturer and scientific advisor to H. Lundbeck A/S. L. Bierut, A. Goate, J. Rice, J.-C.Wang, and the spouse of N. Saccone are listed as inventors on Issued US Patent 8080,371, “Markers for Addiction” covering the use of certain SNPs in determining the diagnosis, prognosis, and treatment of addiction. N. Wodarz has received funding from the German Research Foundation (DFG) and Federal Ministry of Education and Research Germany (BMBF); he has received speaker’s honoraria and travel funds from Janssen-Cilag, Mundipharma, and Indivior. He took part in industry-sponsored multicenter randomized trials by D&A Pharma and Lundbeck. M. Ridinger received compensation from Lundbeck Switzerland and Lundbeck institute for advisory boards and expert meetings, and from Lundbeck and Lilly Suisse for workshops and presentations. K. Mann received speaker fees from Janssen Cilag. H. Kranzler has been an advisory board member, consultant, or continuing medical education speaker for Indivior, Lundbeck, and Otsuka. He is a member of the American Society of Clinical Psychopharmacology’s Alcohol Clinical Trials Initiative, which was sponsored in the past three years by AbbVie, Alkermes, Amygdala Neurosciences, Arbor Pharmaceuticals, Ethypharm, Indivior, Lilly, Lundbeck, Otsuka, and Pfizer. H. Kranzler and J. Gelernter are named as inventors on PCT patent application #15/878,640, entitled “Genotype-guided dosing of opioid agonists,” filed 24 January 2018. J. MacKillop is a principal in BEAM Diagnostics, Inc. D.-S. Choi is a scientific advisory member of Peptron Inc. M. Frye has received grant support from Assurex Health, Mayo Foundation, Myriad, NIAAA, National Institute of Mental Health (NIMH), and Pfizer; he has been a consultant for Intra-Cellular Therapies, Inc., Janssen, Mitsubishi Tanabe Pharma Corporation, Myriad, Neuralstem Inc., Otsuka American Pharmaceutical, Sunovion, and Teva Pharmaceuticals. H. de Wit has received support from Insys Therapeutics and Indivior for studies unrelated to this project, and she has consulted for Marinus and Jazz Pharmaceuticals, also unrelated to this project. T. Wall has previously received funds from ABMRF. J. Nurnberger is an investigator for Janssen. M. Nöthen has received honoraria from the Lundbeck Foundation and the Robert Bosch Stiftung for membership on advisory boards. N. Scherbaum has received honoraria from Abbvie, Sanofi-Aventis, Reckitt Benckiser, Indivior, Lundbeck, and Janssen-Cilag for advisory board membership and the preparation of lectures, manuscripts, and educational materials. Since 2013, N. Scherbaum has also participated in clinical trials financed by Reckitt Benckiser and Indivior. W. Gäbel has received symposia support from Janssen-Cilag GmbH, Neuss, Lilly Deutschland GmbH, Bad Homburg, and Servier, Munich, and is a member of the Faculty of the Lundbeck International Neuroscience Foundation (LINF), Denmark. J. Kaprio has provided consultations on nicotine dependence for Pfizer (Finland) 2012–2015. In the past three years, L. Degenhardt has received investigator-initiated untied educational grants for studies of opioid medications in Australia from Indivior, Mundipharma, and Seqirus. B. Neale is a member of the scientific advisory board for Deep Genomics and has consulted for Camp4 Therapeutics Corporation, Merck & Co., and Avanir Pharmaceuticals, Inc. A. Agrawal previously received peer-reviewed funding and travel reimbursement from ABMRF for unrelated research. All other authors have no conflicts of interest, relevant to the contents of this paper, to disclose.

## Authors Contribution

M. Munn-Chernoff, C. Bulik, and A. Agrawal were responsible for the study concept and design. M. Munn-Chernoff, E.C. Johnson, and Y.-L. Duan performed the statistical analyses, and J. Coleman, R. Walters, and Z. Yilmaz assisted with the data analysis. M. Munn-Chernoff, E.C. Johnson, Y.L. Duan, J. Coleman, L. Thornton, R. Walters, Z. Yilmaz, J. Baker, C. Hübel, J. Kaprio, H. Edenberg, C. Bulik, and A. Agrawal assisted with interpretation of findings. H. Kranzler, J. Gelernter, and H. Zhou facilitated access to and interpretation of the Million Veteran Program summary statistics for AUD. M. Munn-Chernoff, E.C. Johnson, L. Thornton, C. Bulik, and A. Agrawal drafted the manuscript. All remaining authors provided data for this study and consulted on the analytic plan. All authors critically reviewed the content and approved the final version for publication.

## Footnotes

1 In Liu et al. (2019), the phenotype is noted as “smoking cessation”, where current smokers were coded as 2 and former smokers were coded as 1. Because the comparison group is “current smokers”, we have renamed this phenotype as “current smoking” for clarification and ease of interpretation across all smoking phenotypes.

## Notes

https://www.med.unc.edu/pgc/results-and-downloads/

https://conservancy.umn.edu/handle/11299/201564

https://ipsych.au.dk/downloads/

